# Habitat-specific temporal variation in the pace of fish diversification

**DOI:** 10.64898/2026.02.02.703334

**Authors:** Nick Peoples, Michalis Mihalitsis, Peter C. Wainwright

## Abstract

Speciation and extinction events are concentrated unevenly through space and time, shaping the global distribution of extant biodiversity. Habitats modulate these dynamics over short timescales by determining the ecological landscape and providing substrate for diversification. This raises questions about whether biotic and abiotic heterogeneity across habitats also influenced diversification rate over geological timescales. Here, we show the rate of species diversification varies through time according to major differences between aquatic habitats for ray-finned fishes, which comprise over half of vertebrate diversity. Phylogeny-wide net diversification rates have accelerated 1.5-1.7x in reef-associated, freshwater benthopelagic and freshwater demersal lineages, which inhabit complex benthic habitats, while remaining constant through time in other less complex habitats such as the pelagic zone. Combined evidence across multiple diversification models implicates the rise of benthic-feeding fishes, which capitalized on functional feeding innovations and the reorganization of benthic resources following the Paleocene-Eocene Thermal Maximum. Our results show that the global biodiversity of ray-finned fishes is shaped by a combination of clade-specific pulses and tree-wide rate shifts, an outcome of the dynamic interplay between the traits of species and the features of the habitats they occupy.

**Significance statement:** Aquatic habitats vary considerably in their biotic and abiotic properties. In this study, we demonstrate that these differences have modulated the pace of species diversification through time for ray-finned fishes, a group that represents over half of extant vertebrate species. By estimating habitat-specific speciation and extinction rates at a high temporal resolution, we find accelerated diversification beginning ∼50 Mya for species living in complex benthic habitats. We show that this pattern is in part driven by the rise of species that took advantage of functional innovations to feed on a diverse array of benthic associated resources, like coral and algae. These results highlight the interplay between the traits of species and the properties of the habitats they occupy in generating the biodiversity patterns observed today.

## Introduction

Diversity is one of the main currencies of evolutionary biology. Its oscillating dynamics reflect speciation and extinction events concentrated unevenly through space and time^1–7^, yet debate remains over the underlying causes of this temporal variation. Periods of increased diversification may require environmental upheaval or climatic events that reshape the ecological landscape^1,8,9^, organismal modifications that increase the adaptive potential of lineages (e.g., whole genome duplications, functional innovations)^10,11^, or a confluence of the two^12^. However, studies more often find that diversification slows through time, an emergent pattern across many groups^7,13–16^. These temporal slowdowns are frequently attributed to the existence of ecological limits – the accessibility of open niches, resources, or geographic area – which generate diversity-dependent patterns as ecological space becomes saturated^13,16–19^. Species diversity often scales with these properties (e.g., geographic area)^20–22^, yet whether and how these ecological factors modulate the rate of species diversification over long timescales is much less studied. Associating these ecological differences with varied diversification rate trends can provide crucial insight into how and why diversification rates change through time. A species’ habitat is a multivariate representation of biotic and abiotic characteristics that together provide the physical substrate for diversification to occur. Habitats vary considerably in their range of microhabitat gradients, trophic niches available, stability, and connectivity, and this heterogeneity has been shown to shape speciation dynamics over short timescales^23–27^. This raises the possibility that the distribution of extant species diversity has been generated by the accumulated effects of habitat-specific variation in the pace of diversification through time.

Over 35,000 species of ray-finned fishes (Actinopterygii), representing over half of extant vertebrate species, have come to dominate aquatic habitats worldwide. These habitats vary considerably in ecological resources, space, stability and thus, they also vary in the potential for diversification rates to change as opportunity is extinguished or environmental change occurs. The heterogeneity of complex benthic habitats such as shallow coral reefs fosters interactions with multiple, highly diverse resources, promoting niche differentiation and supporting higher species diversity^25,28–30^. Freshwater systems have continuous habitat gradients and abundant physical barriers to gene flow that stimulate divergence events^24^. In contrast, environments with comparatively greater homogeneity such as the pelagic zone and deep sea have narrowed ecological opportunities and high connectivity, decreasing the chances of speciation events^31–34^. These habitats are more stable over geological timescales^35,36^. Actinopterygian diversity is also characterized by remarkable contrasts; freshwater and marine species richness is similar despite vastly greater habitat area in the oceans^37,38^. This suggests that temporal trends in the pace of diversification may vary between major aquatic habitats, affording an opportunity to understand how past diversification dynamics have shaped contemporary patterns of biodiversity. Here, we estimate habitat-specific temporal trends in diversification rates of ray-finned fishes at a global scale. We estimate these rates at an unprecedented resolution to address how major ecological differences (e.g., geographic area, stability, structural complexity) between habitats influence the rate at which species accumulate through time. We compare diversification rate trends among seven major aquatic habitats, identifying both tree-wide and clade-specific rate shifts across ray-finned fishes and associate these trends with shared habitat traits. By incorporating multiple lines of evidence, we show that accelerating diversification rates are a unique attribute of complex benthic habitats in both marine and freshwater systems, a trend fueled by the emergence of clades that capitalized on the diverse array of benthic resources following major ecological stimulation.

## Results and Discussion

### Diversification rates through time

To investigate how diversification rates have changed throughout the evolutionary history of ray-finned fishes, we classified 9,560 species into seven discrete categories (reef-associated, marine demersal, marine benthopelagic, marine pelagic, freshwater demersal, freshwater benthopelagic, freshwater pelagic) representing major aquatic habitats that vary in the ecological opportunities they offer lineages^25,28–30^. We estimated speciation and extinction rates in 1-My intervals for lineages within each habitat using episodic birth-death (EBD) models implemented in RevBayes^39,40^. Ray-finned fishes show substantial variation in the tempo of diversification through time across habitats (Fig. 1a). Net diversification rates are remarkably constant in marine pelagic (MP), marine demersal (MD), marine benthopelagic (MB), and freshwater pelagic (FP) lineages, which inhabit comparatively homogenous environments that represent the majority of aquatic habitat availability by area and volume (Fig. 1a)^41^. This temporal consistency may be a consequence of both the stability of these habitats over long timescales and/or the lack of substrate they provide to stimulate species diversification along multiple ecological axes^31,35^. In contrast, reef-associated (RA) lineages and those in freshwater demersal (FD) and freshwater benthopelagic (FB) habitats show a notable increase in net diversification rate beginning in the Eocene (56-33.9 Ma) and continuing until the Middle to Late Miocene (16-5.3 Ma) (Fig. 1a).

**Figure 1.**
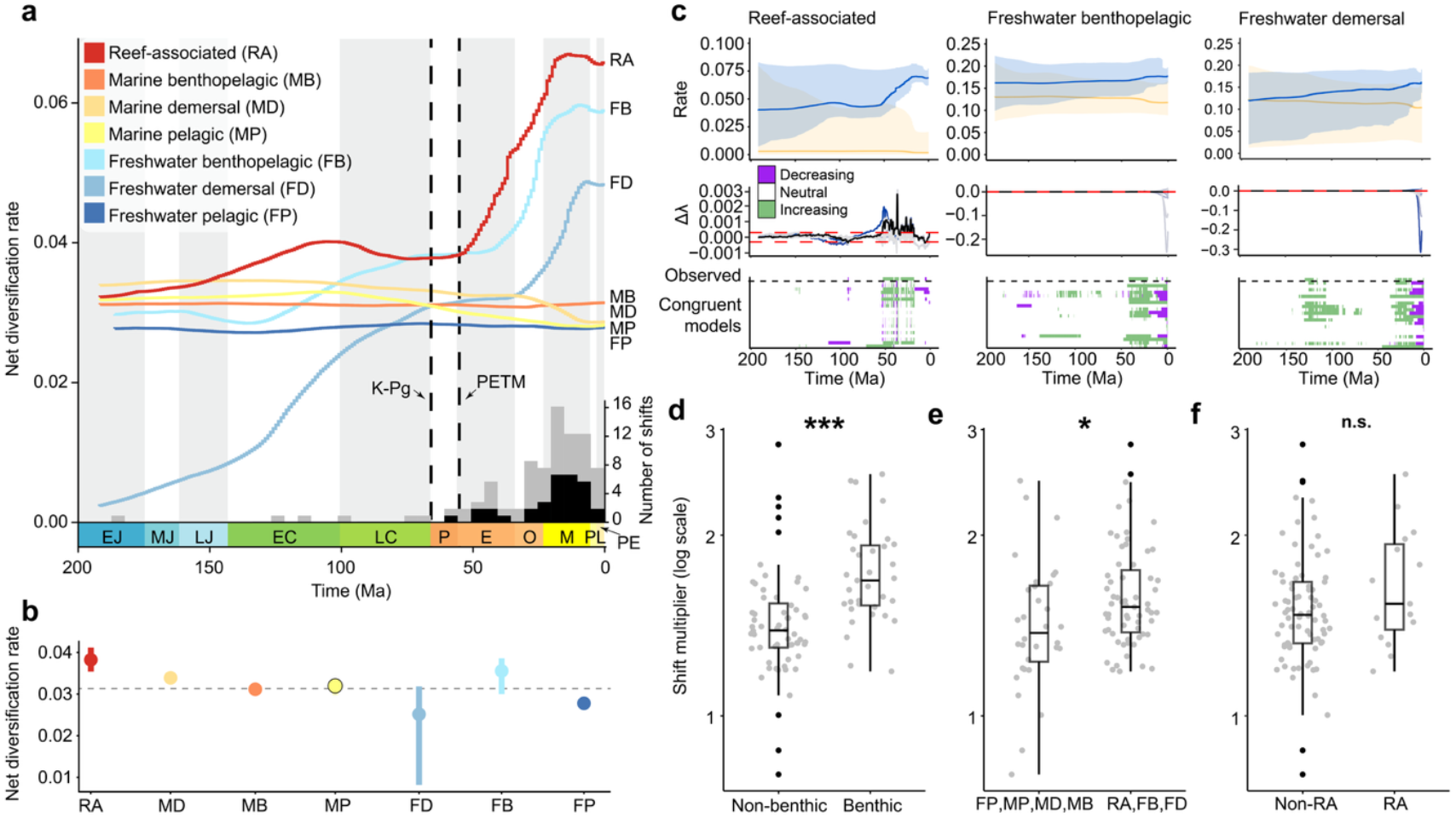
The tempo of ray-finned fish diversification through time. **a**, Net diversification rates in 1-My intervals for reef-associated (RA; n=1907), freshwater benthopelagic (FB; n=3107), freshwater demersal (FD; n=1647), marine benthopelagic (MB; n=300), marine demersal (MD; n=1625), marine pelagic (MP; n=635) and freshwater pelagic (FP; n=339) lineages. Abbreviations of geological epochs: EJ, Early Jurassic; MJ, Middle Jurassic; LJ, Late Jurassic; EC, Early Cretaceous; LC, Late Cretaceous; P, Paleocene; E, Eocene; O, Oligocene; M, Miocene; PL, Pliocene; PE, Pleistocene. Stacked bar chart reporting the number of branch-specific rate shifts through time; black bars correspond to primarily benthic feeding clades. **b**, Median rates for each habitat; the dashed line represents the global median. Bars represent the interquartile range. **c**, Speciation (blue) and extinction (orange) rates, with shaded areas representing 95% confidence intervals, and the range of speciation rate trends under congruent diversification scenarios for RA, FB, and FD lineages. Qualitative trends are reported relative to the observed data. The magnitude of branch-specific rate shifts between **d**, benthic (n=38) and non-benthic (n=57) feeding clades (*P*<0.001), **e**, habitats undergoing net diversification rate increases through time (n=63) and those that do not (n=32) (*P*<0.05), and **f**, reef (n=15) and non-reef associated (n=80) taxa (*P*=0.168). Black points represent outliers.

During this period, tree-wide net diversification increased 1.5-1.7x, after which rates stagnated and gradually decreased towards the present. The rising net diversification of RA, FB, and FD lineages is thus a relatively recent trend in the evolution of ray-finned fishes; median rates vary only slightly (0.025-0.038 lineages/My^-1^) over the last ∼200 My (Fig. 1a). While increasing speciation (λ) explains this trend in RA lineages, declining extinction (μ) underlies net diversification increases in FD and FB lineages (Fig. 1c; *SI Appendix*, Fig. S1). These temporal trends indicate that despite accounting for a minority of total area^41,42^, these habitats (RA, FD, FB) may have shared properties that promote the continued and accelerated generation of species diversity. These habitats are considered more heterogenous, complex, and vulnerable to change^24,28,29^ indicating that the temporal trends of ray-finned fish diversification (Fig. 1a-c) may reflect key differences in the potential for diversification that these aquatic habitats provide.

Given that speciation and extinction rates estimated under birth-death models are not identifiable^43^, we verified that these patterns of diversification rate change between time intervals are consistent under alternative diversification scenarios^44,45^. By jointly sampling alternative λ and μ rate functions to create congruent diversification scenarios, we find consistent patterns of speciation rate change between ∼50 and ∼10 Ma that closely match those under the observed EBD model (Fig. 1c; *SI Appendix*, Fig. S8). We also investigated whether increasing net diversification rates during this same time period can occur under four specific alternative diversification scenarios: constant λ, constant μ, exponentially decreasing λ, and exponentially increasing μ. We find that the observed net diversification rate trends in RA, FD, and FB lineages are either valid under these four scenarios, or the scenario is not contained within the congruence class (e.g., λ or μ < 0 at any point in time) (*SI Appendix*, Figs. S4-S6)^44,45^. In our implementation of the Horseshoe Markov random field (HSMRF) birth-death model, the total rate variability through time is determined in part by the global scale hyperprior (ζ)^40^. We confirmed that the temporal diversification patterns are robust under more “permissive” priors, which effectively allow for more total variation in λ and μ across all intervals (see Materials and Methods; *SI Appendix*, Fig. S2).

While tree-wide shifts in diversification rate may be a consequence of processes affecting clades equally across a broad phylogenetic scale (e.g., climatic events occurring at a global scale), identifying the specific clades that drive these shifts and their commonalities can provide ecologically-centered explanations for why diversification rates change. We used a lineage-specific birth-death-shift (BDS) model^46^ implemented in RevBayes to estimate the timing, magnitude, and phylogenetic placement of speciation rate shifts. We identified 95 rate shifts (posterior probability [PP] of shift > 0.5) which were distributed relatively evenly across habitats when accounting for tree size (RA; 0.0039 shifts per branch, FB; 0.0042, FD; 0.0067, MD; 0.004, MB; 0.005, MP; 0.0071, FP; 0.0104). However, these shifts were distributed unevenly through time and peaked during the Early Miocene (23-16 Ma), a time of considerable climatic and ecosystem reorganization globally (Fig. 1a). The magnitude of rate shifts is higher in RA, FB, and FD lineages (i.e., habitats in which diversification increases through time) (Analysis of Variance [ANOVA]; F=5.904, *P*<0.05) (Fig. 1e), indicating that the emergence of some clades within these habitats (e.g., Pomacentridae, Holocentridae, Acanthuridae [RA], Cichlidae, Cyprinidae [FB], Loricariidae, Sicydiinae, Balitoridae [FD]) may have had a disproportionately larger impact on the tempo of tree-wide net diversification. Though recent net diversification rates are highest in reef taxa (0.066 lineages My^-1^), reef-associated lineages do not have larger shifts than other habitats (ANOVA; F=1.935, *P*=0.168) (Fig. 1f), suggesting an interaction between patterns emerging across the phylogeny (e.g., tree-wide shifts for all lineages within a habitat) and clade-specific rate shifts within these habitats.

Association with the benthos has recently been appreciated as a significant driver of diversification in fishes^32,47,48^, so we compared the magnitude of rate shifts among clades that do and do not feed on benthic-associated resources. We found that primarily benthic-feeding groups undergo 1.2x larger shifts (ANOVA; F=19.73, *P*<0.001) (Fig. 1d) and comprise 40% of all shifts across fishes (Fig. 1a). Our results therefore suggest that the emergence of clades that evolved to successfully exploit benthic resources, coupled with complex habitats and the plethora of ecological niches they offer, are primary contributors to the uneven diversity of extant ray-finned fishes.

### Benthic feeding on reef ecosystems

Change in the pace of diversification is a signal of evolutionary radiation^49^. While slowdowns suggest extinguishing ecological opportunity, increasing rates may imply that environmental change or organismal innovation broadened the range of achievable niches. Marine reef ecosystems are one of the most complex aquatic habitats and long-term cradles of biodiversity^50,51^. Our results show that significant speciation rate shifts in reef lineages most often occur in benthic-feeding groups (60% of shifts within reef-associated lineages), resulting in the largest and fastest increase in net diversification since the Cretaceous-Paleogene (K-Pg) extinction event (Fig. 1a), which raises the possibility that benthic feeding stimulated these changes. While several reef-associated clades have undergone diversification rate shifts, separating the effect of reef habitat itself from other factors such as organismal traits has remained difficult^52–54^. Therefore, we further examined how transitions to feeding on benthic-associated resources shaped the tempo of reef fish diversification.

Using a modified classification of reef fish feeding modes^47^ to reflect whether primary diet items either require detachment (e.g., algae) or are firmly associated with the reef benthos (e.g., urchins), as opposed to free-moving (e.g., plankton), we fit a state-dependent speciation and extinction (SSE) model with hidden states^55^ in RevBayes to estimate speciation and extinction rates of lineages feeding on benthic-associated and non-benthic-associated resources while accounting for background rate variation. Net diversification rates are 1.33-1.57x higher (*T*_*1*.*3A*_=0.909; *T*_*1*.*3B*_=0.592; see Materials and Methods) when lineages transition to feeding on benthic-associated resources, such as coral or algae (Fig. 2c). This effect of benthic feeding on net diversification is robust to an alternative prior distribution of extinction rates (*SI Appendix*, Fig. S7). Though rates estimated under SSE models are not time-varying, examining the distribution of hidden states through time can reveal temporal diversification patterns because positive rate shifts can be reflected by transitions to hidden states with a higher background diversification rate. The proportional rise of benthic feeding fishes in hidden state B (in which net diversification rates are 3.35x higher than hidden state A) since the Paleocene-Eocene Thermal Maximum (PETM) mirrors the tree-wide increase in net diversification rate observed under the EBD model (Fig. 1a, 2b). The acceleration of diversification rate in reef-associated lineages is thus for the most part driven by the rise of butterflyfishes (Chaetodontidae), angelfishes (Pomacanthidae), surgeonfishes (Acanthuridae), triggerfishes (Balistidae), rabbitfishes (Siganidae), parrotfishes (*Scarus, Chlororus*), and damselfishes (Pomacentridae) (Fig. 2a, 3), which all together dominate the benthic-feeding fauna on modern reefs. The emergence of specialized benthic feeders that exploited the habitat complexity and abundance of food resources on reefs is thus a major contributor to the acceleration of species diversification over time in reef associated lineages, which represent a large evolutionary radiation whose diversification has only recently stalled (Fig. 1a).

**Figure 2.**
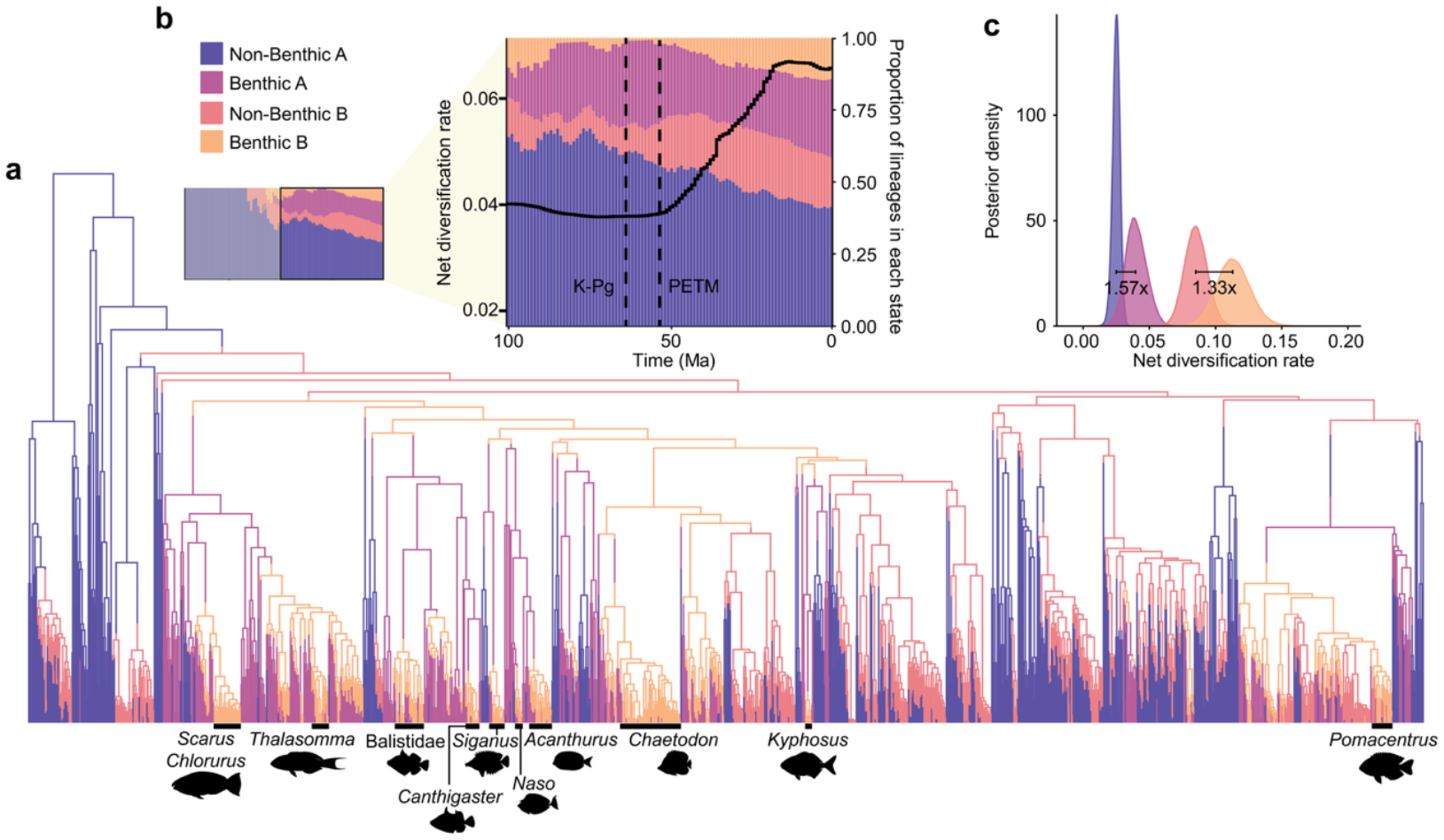
The evolution of benthic-feeding reef fishes. **a**, Ancestral state reconstruction using the HiSSE-2 model of benthic feeding on a phylogeny of coral reef fishes. **b**, The proportion of benthic and non-benthic feeding across two hidden states (A & B) through time, in 1-My intervals. The trend line corresponds to the net diversification rates for reef-associated lineages estimated over the same interval. **c**, Posterior distribution of net diversification rates for benthic and non-benthic feeding lineages across two hidden states. The difference between median rates of feeding types (benthic vs. non-benthic) within each hidden state is reported.

**Figure 3.**
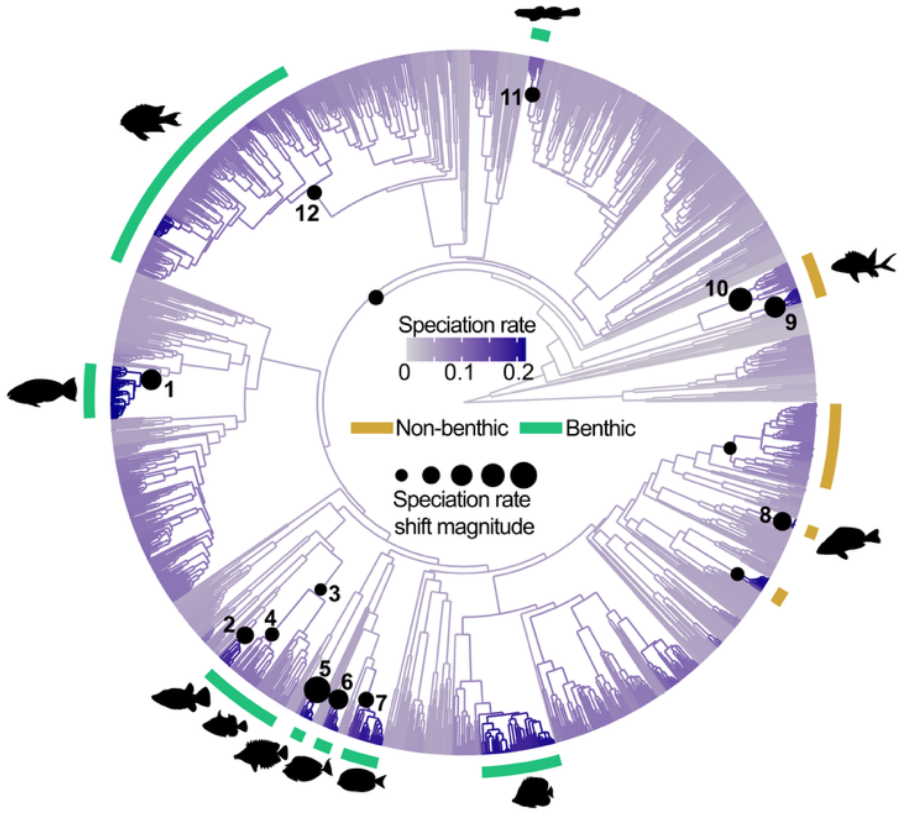
Shifts in the tempo of reef fish diversification. Phylogeny of reef-associated fishes, with black points indicating nodes where significant rate shifts were detected; points are scaled to the magnitude of speciation rate change between parent and daughter branches. Speciation rate shifts in 1) *Scarus* and *Chlororus*, 2) *Canthigaster*, 3) Balistidae, 4) *Rhinecanthus*, 5) Siganidae, 6) *Naso*, 7) *Acanthurus*, 8) *Hypoplectrus*, 9) and 10) Holocentridae, 11) *Elacatinus*, 12) Pomacentridae. Green bars represent predominantly benthic-feeding groups. Ectoparasite feeding (Elacatinus) requires detachment of resources and thus, functionally resembles a biting-associated feeding mode rather than a suction-based mode.

Evolutionary processes operating across broad spatial and temporal scales could generate consistent diversification trends across phylogenetically-disparate groups^56^. In this case, we would expect to find net diversification rate increases over a similar time period in other dominant reef fauna. Major lineages of reef-building scleractinian corals emerged during the Cretaceous and Paleogene, resulting in a structural overhaul of reef habitat complexity^57–59^. We therefore tested if reef-associated corals showed a similar temporal trend in diversification rates as reef-associated fishes. Net diversification rates estimated for 842 species of reef-associated scleractinian corals are remarkably constant through time periods during which large changes are seen in reef fish diversification (but see ref. ^59^) (*SI Appendix*, Fig. S3). This contrasts the pattern observed in reef-associated fishes, suggesting that the increased diversification during this same period is thus likely linked to the evolution of fishes themselves and their ecological interactions, rather than reflecting a general pattern of accelerated diversification across reef-associated taxa.

### Changing ecological opportunities and innovations

Geological and climatic events occurring globally over evolutionary timescales can stimulate major ecological reorganization, fundamentally changing the nature of how lineages interact with each other and their environment^60,61^. The abrupt increase in net diversification rate for RA, FD, and FB lineages occurred during a time of substantial change in aquatic systems and their faunal compositions, marked by the diversification of distinct acanthomorph families and shifts towards low trophic level feeding^62–64^. Intense warming during the PETM ∼56 Ma massively reorganized aquatic food webs and habitats^65–69^ by transforming the benthic resources available to fishes, such as algae and coral^70,71^. The unequal acceleration of species diversification across habitats following this hyperthermal event (Fig. 1a) suggests the PETM and its residual effects may have disproportionately affected freshwater and shallow marine environments by changing the composition of benthic resources, revitalizing opportunity to diversify along novel ecological axes. Some benthic communities persisted through this hyperthermal event due to thermal tolerance (e.g., molluscs^72^), providing both stability and turnover of benthic resources. Concordant mass extinction of established lineages during or immediately following the PETM may have promoted further diversification by reducing competitors^73^, though diverse fish assemblages persisted through this time^74^. In contrast, stable environments with lower ecological opportunity such as the open ocean may have experienced reduced PETM impacts^35^, leading to the stagnation of species diversification (Fig. 1a) in the absence of major upheaval. The steady pace of diversification through time in these habitats suggests that without input of new ecological substrate to stimulate diversification, dominant lineages likely emerged without widespread diversification.

While extensive restructuring of ecological resources in freshwater and marine reef systems may have opened ecological opportunity, lineages are not equally equipped to exploit these changes. Feeding on benthic resources involves various forms of biting mechanics, which strongly contrasts the ubiquity of prey capture by suction feeding on free-moving prey. Multiple innovations in the feeding system characterize the evolution of benthic-feeding fishes. These include advances in both acquisition (e.g. complex multicuspid teeth, lateral jaw movement, intramandibular joint)^75–77^, processing (e.g., pharyngeal mill)^78^, and assimilation^79,80^ which together increase performance in these feeding niches. The effect of these clades on diversification is remarkably consistent across models (episodic birth-death [Fig. 1a, b], birth-death-shift [Fig. 1a, 3], state-dependent speciation and extinction [Fig. 2]), suggesting that benthic-feeding fishes may represent lineages that withstand the loss of speciation potential through time^7^ to generate pulses of increased diversity across the actinopterygian phylogeny. This intertwined interaction between organismal functional innovation and major environmental upheaval most likely caused the recent and revitalized diversification of fishes in complex benthic habitats.

## Conclusion

Aquatic habitats differ in their ability to stimulate diversification, leading to major consequences for the distribution of extant biodiversity patterns. Our results add resolution to a striking paradigm. Although the number of freshwater and marine ray-finned fish species is relatively equal^37^, nearly 70% of species occupy just three complex benthic habitats, a pattern that can be attributed to their accelerated diversification rates. Microhabitat gradients and high structural complexity in freshwater benthic and marine reef habitats provide opportunity in waiting for properly equipped lineages to exploit, whereas vast comparatively homogenous habitats such as the open ocean promote steady diversification through time. Jointly, benthic-feeding lineages that evolved functional feeding innovations capitalized on renewed ecological opportunities following global environmental change. This dynamic interplay between the traits of species and the attributes of benthic habitats prompted the progressive acceleration of species diversification over ∼30 My. The resultant tree-wide rate shifts and clade-specific pulses structured the distribution of contemporary fish biodiversity.

## Materials and methods

### Phylogeny and habitat classification

For all phylogenetic comparative analyses, we used the molecular phylogeny of Rabosky et al.^81^. While other phylogenetic hypotheses for subsets of fishes exist, we opted to use a single time calibration to ensure our analyses are commensurate and comparable across habitat subtrees. We used FishBase^82^ to classify species into seven discrete habitats: marine reef-associated (n=1907 species), marine demersal (n=1625 species), marine benthopelagic (n=300 species), marine pelagic (n=635 species), freshwater demersal (n=1647 species), freshwater benthopelagic (n=3107 species), freshwater pelagic (n=339 species). We created seven individual habitat trees by pruning the full phylogeny to match each set of species within each habitat. We excluded species that were found in both freshwater and marine (brackish) habitats, freshwater and brackish, marine and brackish, or all three. Within each salinity environment, we combined the following habitats: (1) bathydemersal, demersal, (2) pelagic, bathypelagic, pelagic-oceanic, pelagic-neritic. We recognize that our method does not fully account for the evolutionary history of transitions into and occupation of these habitats.

### Diversification rate through time

To estimate diversification rates across habitats through time, we implemented an episodic birth-death (EBD) model in RevBayes v1.2.1^39^. This model assumes rates are constant within time intervals but can change between intervals. We used a Horseshoe Markov random field (HSMRF) for the prior distribution of log-transformed rates, which assumes autocorrelation of rates between adjacent time intervals. We estimated speciation (λ) and extinction (μ) rates in 1-My intervals for each of the seven habitat trees separately. We set unique global scale hyperpriors (ζ) for each tree, which control the overall variability in diversification rates through time, using the ‘*setMRFGGlobalScaleHyperpriorNShifts*’ function in the R package ‘*RevGadgets*’^83^ with a prior number of shifts=log(2) and a shift size of 2 (2-fold change)^40^ (FD: 368 intervals, ζ=0.0003738458; FB and FP: 187 intervals, ζ= 0.0009208335; RA, MP, MD, MB: 193 intervals, ζ= 0.0008787526). We tested the sensitivity of our results to the choice of the global scale hyperprior by repeating all analyses with ζ values that produce both 5 and 10 prior shifts, a more “permissive” approach that effectively increases the total rate variability through time compared to ζ for log(2) shifts, a conservative value suggested by ref.^40^ that follows the framework of ref. ^84^. For these sensitivity analyses, we maintained an effective shift size of 2-fold. We report results from log(2) shifts in the main text because this prior is reasonably conservative but flexible^40^, and results from 5 and 10 shifts in the supplementary information (*SI Appendix*, Fig. S2). We set unique sampling fractions (ρ) for each tree (RA ρ=0.475, MP ρ=0.303, MD ρ=0.255, MB ρ=0.277, FP ρ=0.355, FD ρ=0.347, FB ρ=0.383). We used Markov Chain Monte Carlo (MCMC) to estimate model parameters; each MCMC had two independent runs for 500,000 generations each with sampling every 10 generations and a pre-burn-in of 50,000 generations (tuning interval=200). This burn-in period helps ensure acceptance rates remain within an acceptable range and improves MCMC efficiency. We further removed 25% of the combined runs as burn-in. We used the R package *convenience*^85^ to verify convergence of speciation and extinction rates in the combined runs at a precision level (α) of 0.015; the posterior distributions of both rates converged for all habitats. We used *RevGadgets* to visualize diversification rates through time; we report net diversification (*r*; λ-μ) rates in the main text. For all figures and analyses on the median rates, we retained only rates over the last 193 million years of the freshwater demersal (FD; root age 368 Ma) tree so that all trees had comparable timescales.

### Congruent diversification scenarios

Diversification rates estimated from birth-death models are not identifiable, however shared patterns across congruent birth-death models can be inferred when diversification rates are found to vary through time^43,44,86^. We explored the validity of alternative hypotheses for diversification rates using the R package *CRABS*^45^ for RA, FD, and FB lineages, which showed rate change through time. We tested hypotheses of constant μ, exponentially increasing μ, constant λ, and exponentially decreasing λ (*SI Appendix*, Figs. S4-S6). We accepted that alternate hypotheses were not contained within the congruence class if μ or λ were negative at any point. We summarized trends in net diversification rate (*r*) change across reference and congruent models with a threshold of twice the mean change in λ between time intervals, which was unique to each habitat. We also simultaneously generated 20 congruent sets of λ and μ through joint sampling with HSMRF *p*-trajectories^86^ (σ_MRF_=1, β_1_=0.5, β_2_=0.2) and visualized trends in speciation and extinction rates across models within the congruence class (Fig. 1c; *SI Appendix*, Fig. S8).

### Coral diversification

We estimated shifts in diversification rate through time for reef-associated scleractinian corals, the major clade of reef-building invertebrates. We summarized the distribution of trees from Huang and Roy^87^ into a single maximum a posteriori (MAP) tree using the ‘*mapTree*’ function in RevBayes v1.2.1 and pruned this tree to match reef-associated species (n=842 species). We used the EBD model described above to estimate diversification rates through time in 1-My intervals (351 intervals, ζ= 0.0003717236, ρ=0.545). The MCMC had two independent runs of 1,000,000 generations each with 25% removed as burn-in. We used Kolmogorov-Smirnov (KS) tests to check for convergence of the replicates (α=0.015; *SI Appendix*, Fig. S3) and combined the runs.

### Branch-specific diversification rate estimation

We implemented a branch-specific diversification rate model in RevBayes v1.2.1 to estimate the timing, location, and magnitude of clade and branch rate shifts^46^. Under this model, both extant and extinct lineages can undergo shifts in diversification rate. Speciation rates were drawn from a log-normal distribution discretized into 6 quantiles (*k*=6); the mean was drawn from a log-uniform distribution with bounds (1E-6, 1E2), and standard deviation drawn from an exponential distribution (lambda=1.7). We used the same prior for the distribution of extinction rates, but constrained extinction rates to be equal across all *k*. The number of rate-shifts was drawn from a uniform distribution with bounds (0, 100). Unique sampling fractions (ρ) were set for each tree; these were equal to those of the EBD analysis and unsampled taxa were assumed to be random. We used MCMC to sample speciation and extinction rates, and stochastic branch rates; two independent chains were run for 500,000 generations each (tuning interval=200), sampling every 10 generations. We combined the runs and removed 50% as burn-in; we used the R package *convenience* to verify convergence (α=0.015). To identify the timing and phylogenetic placement of shifts within each habitat, we first identified all branches where the posterior probability (PP) of a shift was above 0.5. We then quantified the shift magnitude as the ratio between the mean speciation rate on the shift branch and parent branch (multiplier). We compared the magnitude of shifts between reef (n=15) and non-reef (n=80), between benthic (n=38) and non-benthic (n=57), and between RA, FD, FB (n=63) and MD, MB, MP, FP (n=32) using one-way Analysis of Variance (ANOVA) tests. We log-transformed the shift multiplier to fulfill assumptions of normality (Shapiro-Wilk test *P*>0.05).

### Benthic feeding classification

To categorize reef fish species as feeding on benthic associated resources, we modified the diet classifications of Corn et al.^47^. We defined benthic resources as those that require detachment from the reef surface (e.g. turf algae, sponges, coral polyps, mollusks) or are firmly associated with the reef benthos and resist detachment (e.g., urchins, some crabs). Species feeding on elusive free-swimming resources in the water column were categorized as non-benthic feeding. We classified species that feed on ectoparasites (e.g., *Elacatinus*) as benthic-feeding because these resources are attached to a substrate and require detachment. All ‘ram biters’^47^ were categorized as feeding on non-benthic resources; classifications can be found in the Supplementary Data. We pruned the marine reef-associated tree to match the 1288 species for which we had data on feeding mode. We also classified clades in which a speciation rate shift occurred on the parent branch in this same scheme using diet data available in FishBase^82^; we made classifications based on which diet items were most prevalent across species in the clade. We did not classify shifts that occurred in deep nodes that included multiple disparate clades (e.g., at the base of Cypriniformes).

### State-dependent diversification

To test if reef species feeding on benthic-associated resources have higher diversification rates than those that do not, we fit a two-state hidden-state speciation and extinction (HiSSE) model^55^ in RevBayes v1.2.1. The framework of the Bayesian implementation of HiSSE explicitly tests a null hypothesis of character-independent diversification, reducing the need for extensive model comparison. If character-independent diversification is supported, the posterior distribution of net diversification rates within a hidden state should largely overlap. We used two hidden states (HiSSE-2) to accommodate rate variation independent of the observed characters. Speciation and extinction rates were drawn from a normal distribution with mean ln(# taxa/2) divided by the age of the tree (ρ=0.321). We set a prior of 50 state transitions across the tree, and an equal-probability of non-benthic feeding across either hidden state as the prior on the root state. To test if our results were robust to assumptions on the extinction rate, we placed a lower bound on the prior distribution of extinction rates^76,88^. Thus, μ = *A* * λ + *δ* where *A* = 0.5 (μ at least half of λ) and *δ* is a random variable that allows extinction to be greater than *A*\*λ. We used MCMC to sample λ, μ, and stochastic character maps; two independent chains were run for 50,000 generations each (tuning interval=100), sampling every generation, with 5,000 generations pre-burn-in. We combined the runs, removed the first 10% as burn-in, and verified convergence using the *convenience* package (α=0.015). We processed the posterior distribution of model parameters using *RevGadgets*.

To further evaluate the probability that benthic-feeding species have higher net diversification than non-benthic-feeding species, we used a simple test statistic, *T*_*ab*_, where *a* is a specified magnitude of difference (e.g. 1.5x) and *b* specifies the hidden state (e.g. A, B). Thus, *T*_*1*.*5A*_ represents the number of generations for which net diversification for state 1 (*r*_1_) is 1.5x greater than the net diversification for state 0 (*r*_0_), divided by the total number of generations, for hidden state A. This is effectively the posterior probability that *r*_1_>*r*_0_ at a user-specified threshold (e.g., 1.5x greater) within a hidden state under a Bayesian HiSSE model.

## Acknowledgements

Funding for this work was provided by the College of Biological Sciences, University of California, Davis. N.P. was supported by a University of California, Davis Dissertation Year Fellowship and the Achievement Rewards for College Scientists (ARCS) Foundation.

## Data availability

All RevBayes scripts and phylogenetic trees can be found on GitHub (https://github.com/npeoples/fish_temporal_diversification_rate/tree/main).

