## Supplementary material for "Habitat-specific temporal variation in the pace of fish diversification": SI Appendix

**This PDF file includes:**

Figures S1 to S8

**Figures**

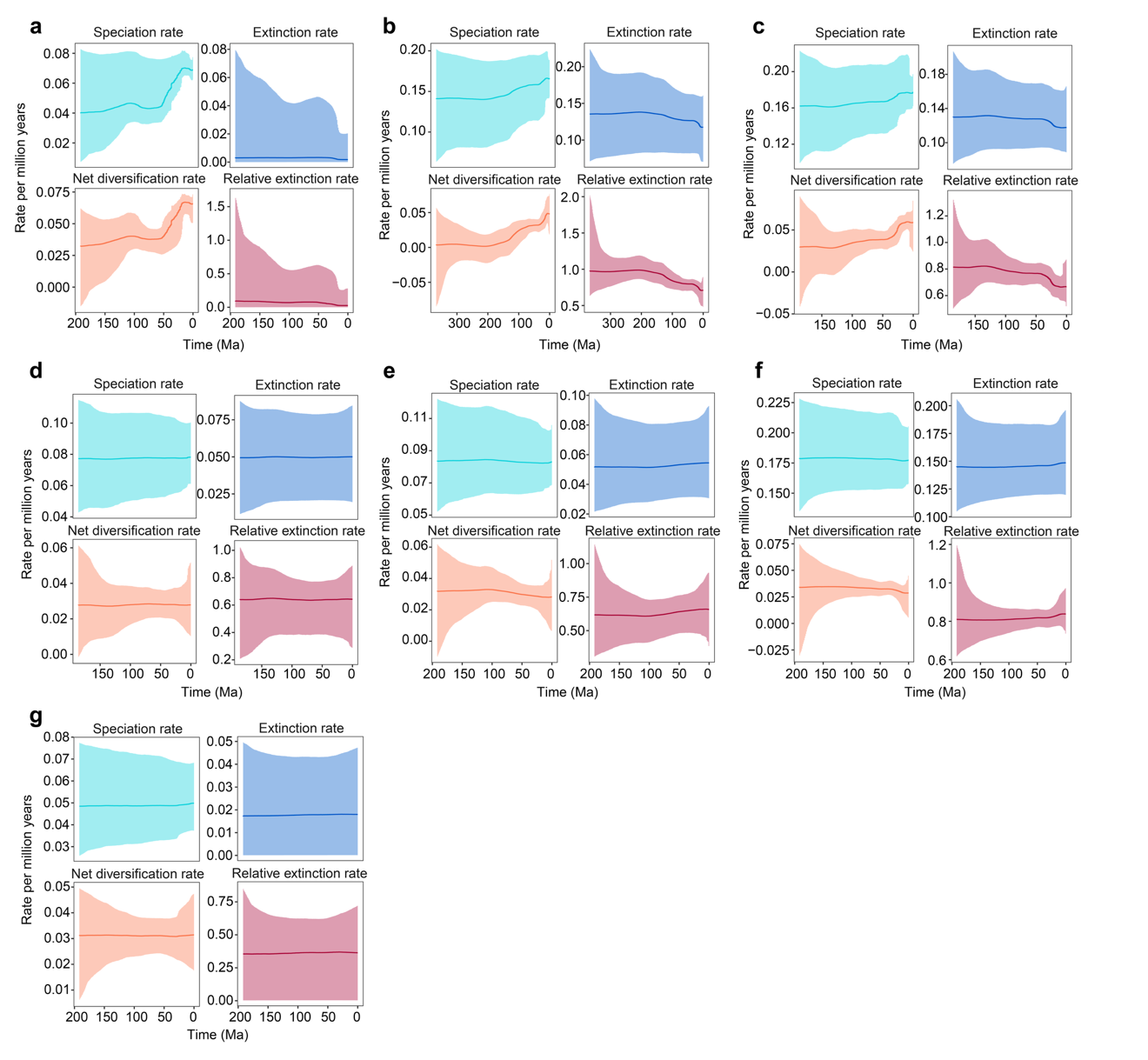

Fig. S1. Speciation, extinction, net diversification, and relative extinction rates through time in 1-My intervals across seven major aquatic habitat types. Shaded areas represent 95% confidence intervals, and relative extinction is reported as the ratio of extinction (μ) to speciation (λ). Rates estimated in 1-My intervals. a, reef-associated (RA), b, freshwater demersal (FD), c, freshwater benthopelagic (FB), d, freshwater pelagic (FP), e, marine pelagic (MP), f, marine demersal (MD), g, marine benthopelagic (MB).

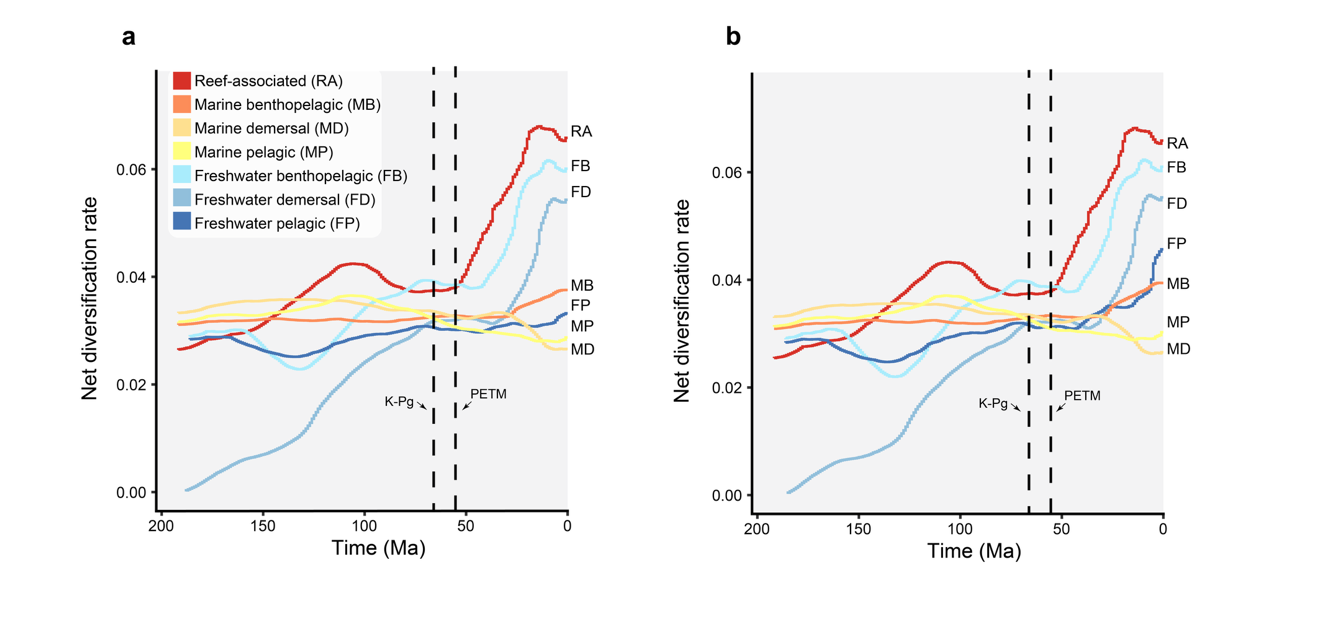

**Fig. S2.** **Temporal trends in net diversification rates across habitats are stable across different global scale hyperpriors.** Net diversification rates estimated in 1-My intervals with a prior of **a,** 5 shifts and **b,** 10 shifts between intervals. The dashed lines mark the K-Pg extinction and the Paleocene-Eocene Thermal Maximum (PETM).

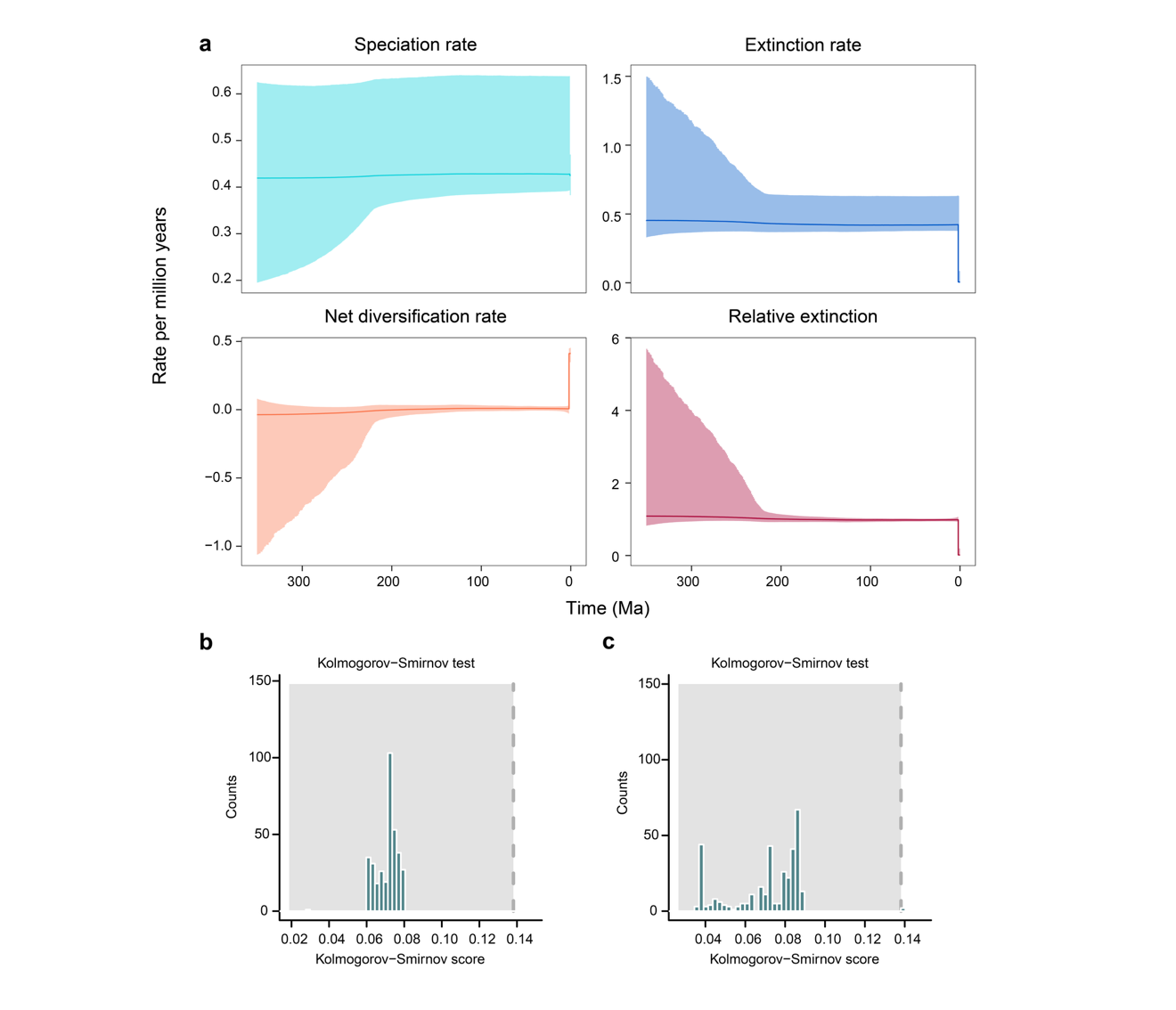

**Fig. S3. Temporal trends in scleractinian (stony coral) diversification rates. a,** Speciation, extinction, net diversification, and relative extinction through time in 1-My intervals for 842 species of reef-associated scleractinian corals. Kolmogorov-Smirnov tests for convergence of **b**, speciation and **c**, extinction rates between runs.

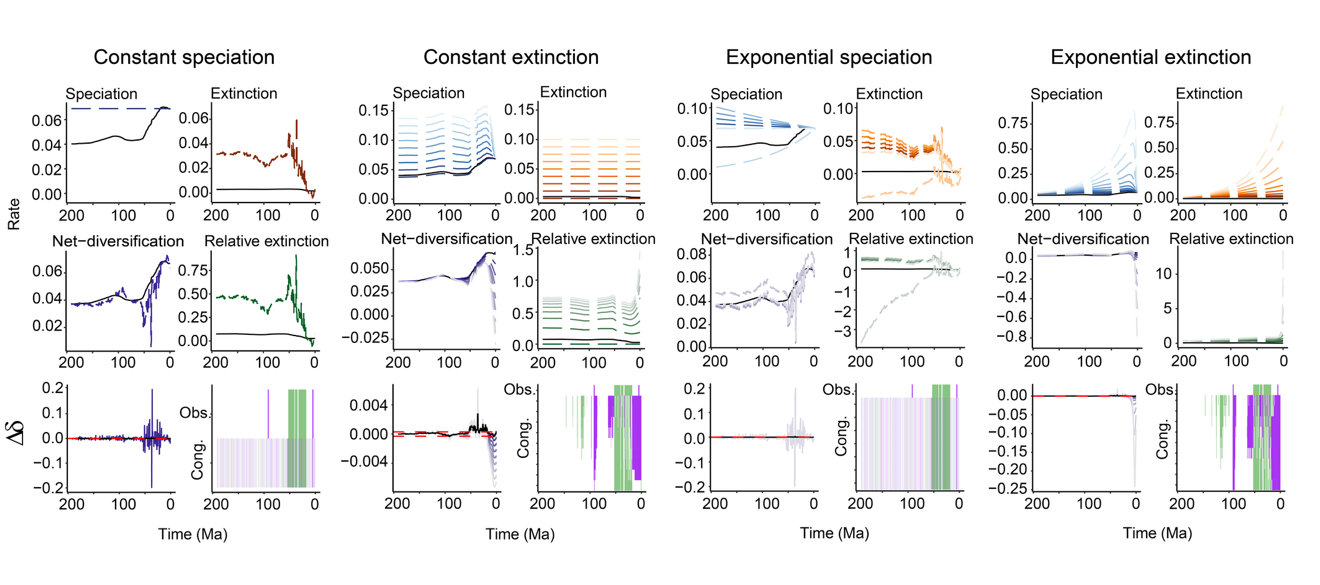

**Fig. S4.** **Congruent diversification scenarios for reef-associated species.** The range of congruent diversification patterns. Constant speciation, constant extinction, exponentially decreasing speciation, and exponentially increasing extinction scenarios are reported. For each scenario, congruent trends are reported relative to the observed rate (solid black line); purple represents qualitative patterns of rate decreases, green represents qualitative patterns of rate increases, and white represents no rate change between intervals.

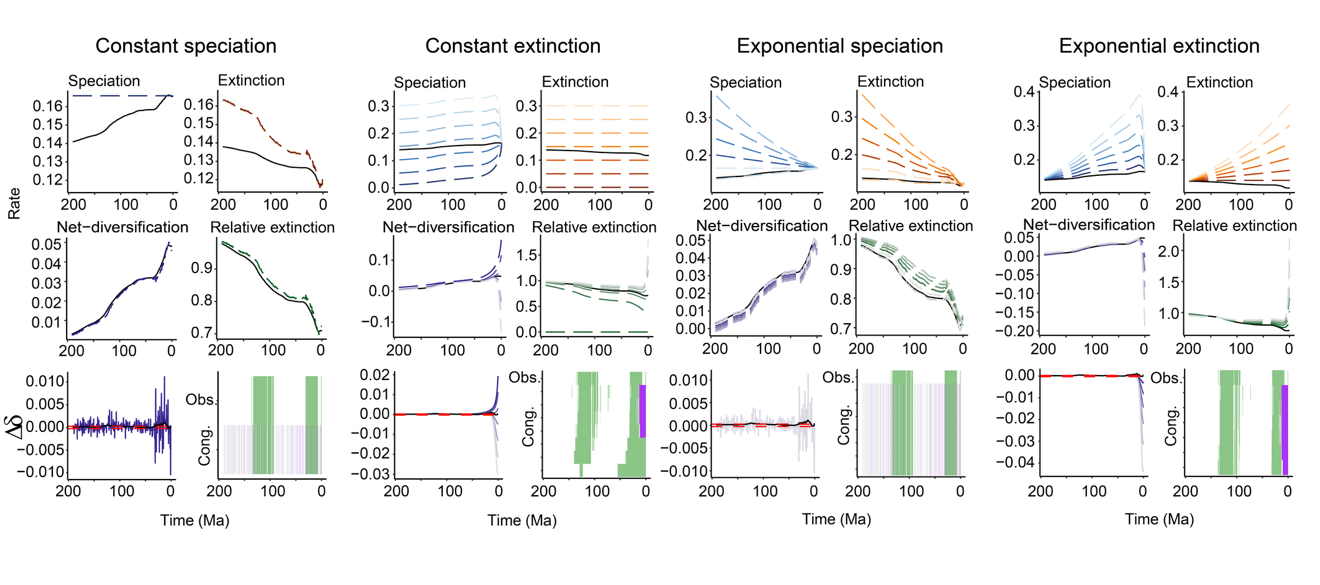

Fig. S5. Congruent diversification scenarios for freshwater demersal species. The range of congruent diversification patterns. Constant speciation, constant extinction, exponentially decreasing speciation, and exponentially increasing extinction scenarios are reported. For each scenario, congruent trends are reported relative to the observed rate (solid blackline); purple represents qualitative patterns of rate decreases, green represents qualitative patterns of rate increases, and white represents no rate change between intervals.

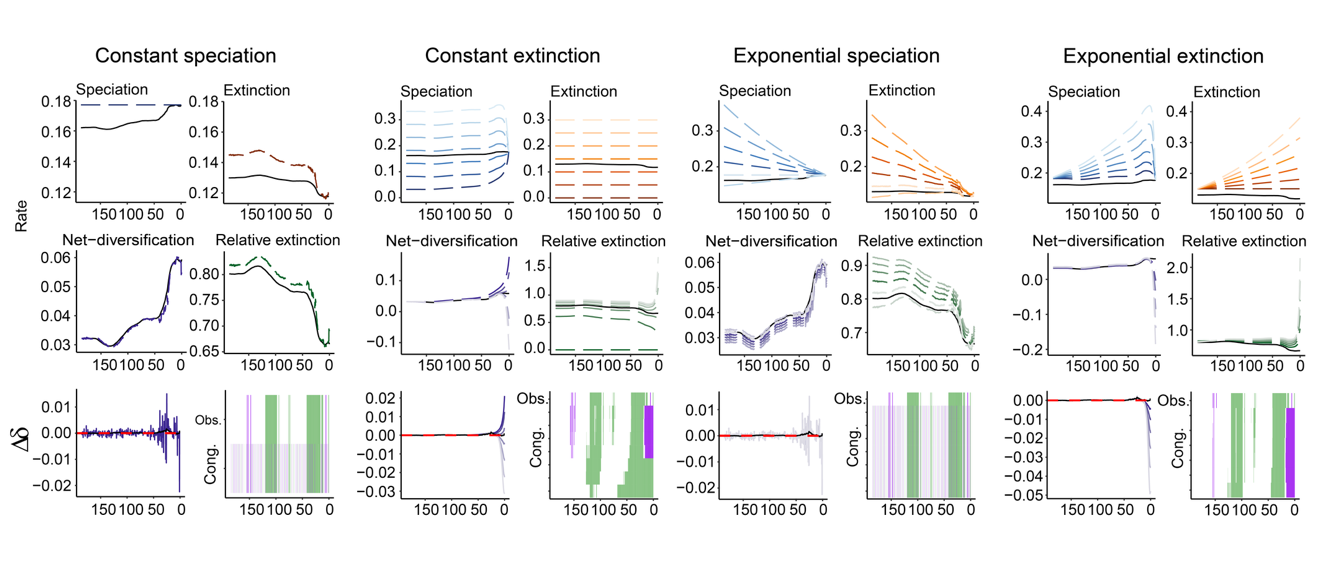

Fig. S6. Congruent diversification scenarios for freshwater benthopelagic species. The range of congruent diversification patterns. Constant speciation, constant extinction, exponentially decreasing speciation, and exponentially increasing extinction scenarios are reported. For each scenario, congruent trends are reported relative to the observed rate (solid black line); purple represents qualitative patterns of rate decreases, green represents qualitative patterns of rate increases, and white represents no rate change between intervals.

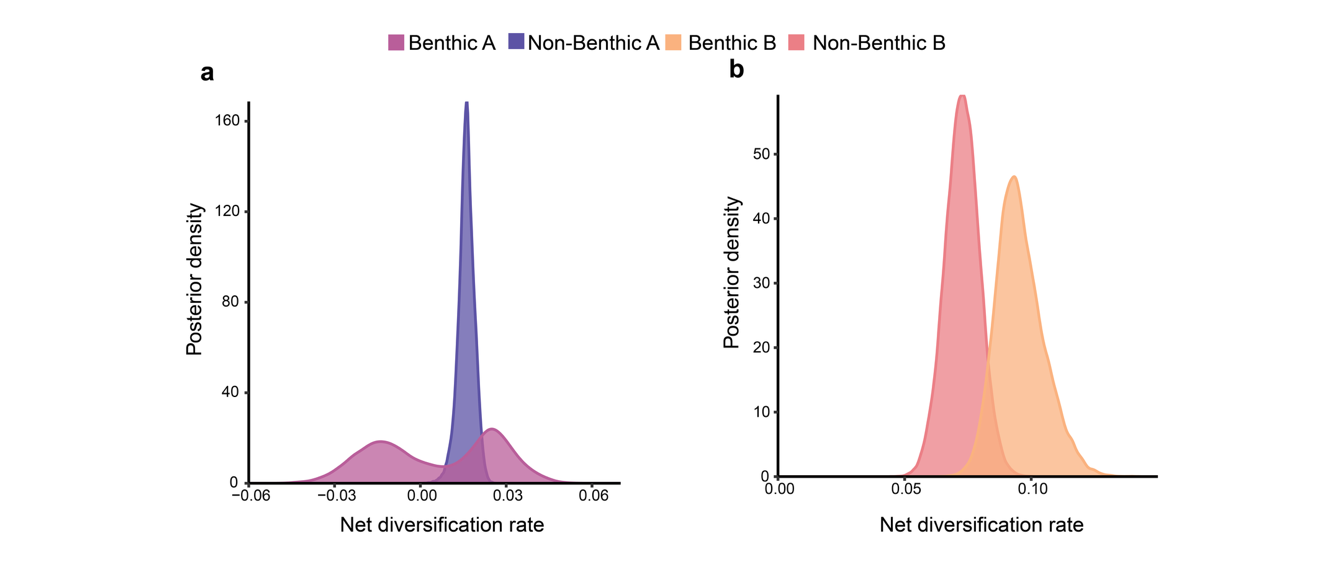

Fig. S7. The effect of benthic feeding is stable under alternate extinction priors. Net diversification rates in a, hidden state A and b, hidden state B, estimated under a HiSSE-2 model, when the extinction rate is at least half of the speciation rate.

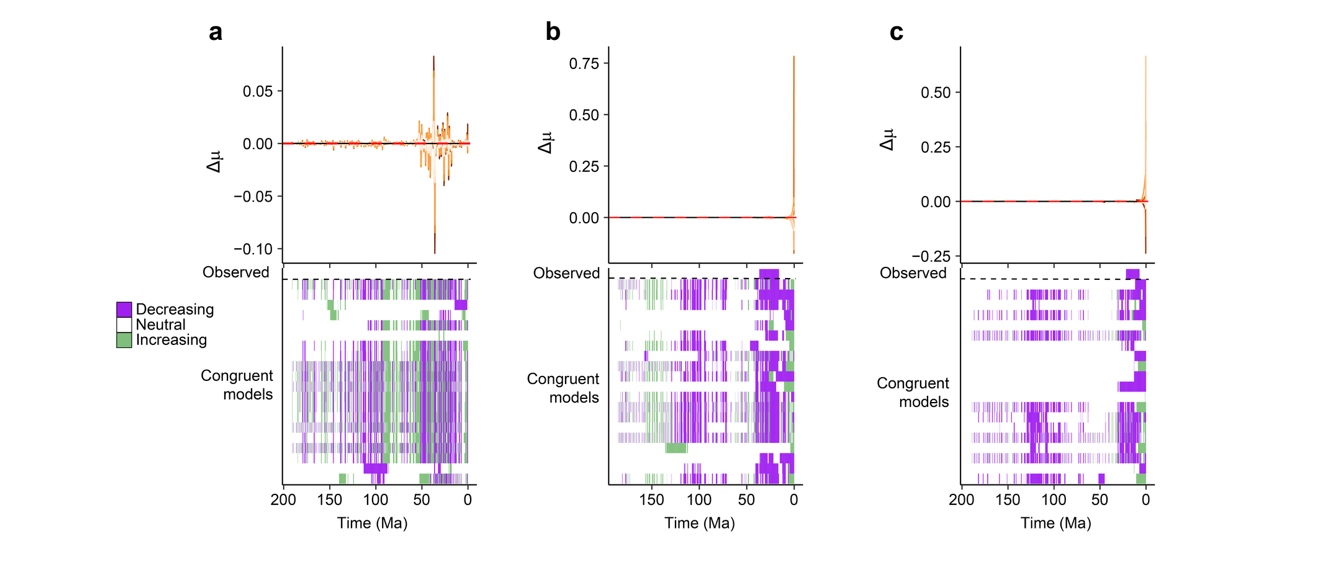

Fig. S8. Extinction rate trends of congruent diversification scenarios. Trends in the extinction rate change between intervals for a, reef-associated lineages, b, freshwater benthopelagic lineages, and c, freshwater demersal lineages under congruent diversification scenarios generated through joint sampling of speciation and extinction rates with horseshoe Markov random field (HSMRF) *p*-trajectories.
